# Rapid evolution of *Wolbachia* in cherry fruit flies

**DOI:** 10.1101/2025.01.13.632697

**Authors:** Daniel J. Bruzzese, Hannes Schuler, Wee L. Yee, Aurel Holzschuh, Jeffrey L. Feder

**Author notes:** Corresponding author, <u>(DJB)</u>. Department of Epidemiology of Microbial Diseases, Yale School of Public Health, Yale University, New Haven, Connecticut, United States of America (Daniel J. Bruzzese).

## Abstract

*Wolbachia* is a widespread bacterial endosymbiont in arthropods known for its ability to manipulate host reproduction through cytoplasmic incompatibility (CI), thereby promoting its spread. However, *Wolbachia* harbors an array of mobile genetic elements (MGEs) that can rapidly alter its genomic structure and content, including the *cif* loci responsible for CI. At the phylogenetic scale, *Wolbachia* genomes are shown to be dynamic with MGEs causing widespread genome rearrangements. But on ecological time scales, the evolution of *Wolbachia* remains largely unknown. In this study, we leverage the natural history of *Wolbachia* in cherry-infesting fruit flies in the *Rhagoletis cingulata* sibling species group. Members of this species group share a common *Wolbachia* strain, *w*Cin2, and their divergence spans from 1,000 to 150,000 years ago. We utilized Nanopore sequencing to characterize *w*Cin2 *Wolbachia* genome divergence across recent to more distant evolutionary timescales. We report rapid evolution of population-level differences in gene content (including *cif* loci and MGEs) and genome structure, with differentiation increasing with time since host divergence. Notably, structural variants were the first to appear both between and among *w*Cin2 populations. Our results also indicated that the CI phenotype previously attributed to a distinct *Wolbachia* strain, *w*Cin3, that coinfects flies in the southwestern USA and Mexico, may instead be caused by *cif* genes that were horizontally transferred into *w*Cin2. Finally, we discovered a novel *Wolbachia* strain, *w*Ind, which appears to have been recently horizontally acquired by cherry fly populations in the Pacific Northwest of the USA. Our findings underscore the fluidity and rapid genome evolution of *Wolbachia*, with significant implications for *cif* gene dynamics and potential impacts on host evolutionary trajectories.

**Author Summary:** In our study, we explore the genome evolution of *Wolbachia*, a common bacterial endosymbiont found in many insects that can manipulate its hosts’ reproduction to help it spread more effectively. *Wolbachia’s* genome is shown to change dramatically over long evolutionary periods due to mobile genetic elements. However, little is known about how *Wolbachia’s* genome evolves over shorter ecological timeframes. Here, we utilized cherry-infesting fruit flies and their *Wolbachia* strain known as *w*Cin2, both of which have spread across North America for the past 150,000 years. Using long-read sequencing, we assembled multiple *w*Cin2 *Wolbachia* genomes from across North America and made comparisons between *w*Cin2 *Wolbachia* genomes that have recently diverged and comparisons to those that have been isolated for over 150,000 years. Our findings reveal that *Wolbachia’s* genome can evolve swiftly, with differences in gene content and most prominently genome structure arising between recently separated *Wolbachia* populations and increasing as populations diverge over time. Interestingly, these changes also impact the key genes responsible for *Wolbachia’s* reproductive manipulation. This research highlights the rapid and dynamic nature of *Wolbachia* genome evolution across different evolutionary timescales and offers insights into endosymbiont-host relationships as well as their potential impact on host speciation and vector control.

## Introduction

The Alphaproteobacteria *Wolbachia* is one of the most common intracellular endosymbionts, infecting over 50% of all terrestrial arthropod species [1], as well as nematodes [2]. Primarily passed vertically from mother to offspring and occasionally horizontally between different host species [3,4], *Wolbachia* teeters between conflict and cooperation with its hosts to aid its spread. In some instances, *Wolbachia* hastens its spread within host populations by positively impacting host fitness via nutrient supplementation [5–7] or disease resistance [8–10]. Conversely, many *Wolbachia* strains selfishly manipulate host reproduction to invade new populations [11].

*Wolbachia’s* reproductive manipulation ranges from male killing, parthenogenesis, feminization, and, most commonly, cytoplasmic incompatibility (CI) [12,13]. CI occurs when male hosts infected with a *Wolbachia* strain mate with uninfected females or females infected with a different, incompatible strain, resulting in the embryonic death of offspring. As a result, selection acts in a frequency dependent manner on CI-inducing *Wolbachia*. Above a critical threshold frequency (usually about 10%), CI-inducing *Wolbachia* can rapidly sweep through a host population [14,15]. Yet, below this threshold, the endosymbiont does not increase in frequency and can be eliminated from host populations [14,15]. CI-inducing *Wolbachia* can potentially contribute to speciation by causing postmating reproductive isolation between hosts infected with different incompatible strains [16–18]. It is also possible that CI can select for increased premating reproductive isolation in its host through reinforcement, by favoring uninfected females that avoid mating with infected males [19,20].

Two *Wolbachia* genes, *cifA* and *cifB* that are associated with the *Wolbachia* prophage WO control the induction and rescue of CI [21–25]. Expression of *cifA* alone in females is sufficient to rescue CI in crosses with infected males [25,26]. The mechanism(s) behind CI induction, however, can vary between strains, with some systems requiring co-expression of *cifA* and *cifB* in male testes to induce CI [13,23,25], while in other systems, the expression of *cifB* alone suffices [27]. Phylogenetic analysis has identified five types (I-V) of *cifA* and *cifB* homologs that have co-diversified [28,29]. Each *cif* type shares common protein domains and the ability to rescue CI if induced by *cif* genes of the same type [13]. Adding further complexity, *cif* gene expression can also be influenced by the genetic background of the endosymbiont host, environmental conditions, and pseudogenization—all of which can impact the strength of CI [30–32].

The dynamic nature of *Wolbachia* genomes plays an important role in mediating both the transfer and expression of *cif* genes. *Wolbachia* genomes contain many active mobile genetic elements (MGEs), including prophages, transposable elements (TEs), and plasmids [33–37]. Active MGEs can transfer both prophage DNA, as well as nearby *cif* gene pairs, between *Wolbachia* strains [28,29,38,39]. In addition to horizontal gene transfer between *Wolbachia* strains, internal rearrangements within the *Wolbachia* genome itself may also alter the expression of *cif* genes. Normally, *cifA* and *cifB* genes are transcribed in tandem, with the *cifA* locus upstream of *cifB* [23]. MGEs can cause translocations or inversions that can separate tandem copies of *cif* genes or insertions/deletions that disrupt *cif* gene expression [40]. Finally, point mutations unrelated to MGEs can also accumulate in *cif* genes and cause their pseudogenization [29,41,42]. Over long-term, phylogenetic scales, *Wolbachia* genomes both accrue genomic rearrangements and exchange their *cif* gene modules, the latter of which is often independent of *Wolbachia* and host phylogenies [39]. There are considerably fewer studies on the short-term evolution of *Wolbachia* genomes on ecological timeframes, however, limited evidence does suggest that *Wolbachia* genomes can evolve rapidly over tens of years [43–45].

Cherry-infesting fruit flies from the *Rhagoletis cingulata* (Diptera: Tephritidae) sibling species group and their associated *Wolbachia* strain, *w*Cin2, provide an excellent system to study *Wolbachia* genome evolution across ecological to phylogenetic evolutionary timescales. *Rhagoletis indifferens* Curran, is endemic to the Pacific Northwest (PNW) region and Northern California of N. America and infests the fruit of bitter cherry (*Prunus emarginata*) [46] (Fig 1A). *Rhagoletis cingulata* (Loew), attacks black cherry (*P. serotina*) in eastern North America (ENA), the southwestern USA (SW), and in the Sierra Madre Oriental Mountains (SMO) and the Eje Volcánico Transversal de Mexicano (EVTM) regions of Mexico (Fig 1A) [47–50]. Genetic surveys of microsatellite loci revealed a pattern of clinal geographic variation among cherry flies across N. America, suggesting populations in the PNW, ENA, SW, SMO, and EVTM became isolated from one another due to the onset of warmer and drier conditions at the end of the Pleistocene and beginning of the Holocene ∼15,079 ya (95% credible interval 7,143 to 31,270 years) (Fig 1B) [49]. In contrast, mtDNA haplotypes display a disjunct geographic divide that differentiates cherry fly populations (Fig 1B). In this regard, *R. cingulata* in the SW, SMO, and EVTM possess a distinct mtDNA haplotype that differs from that shared between *R. indifferens* in the PNW and *R. cingulata* in ENA populations, with an estimated divergence time of ∼100,000 to 157,000 years [50].

**Fig 1.**
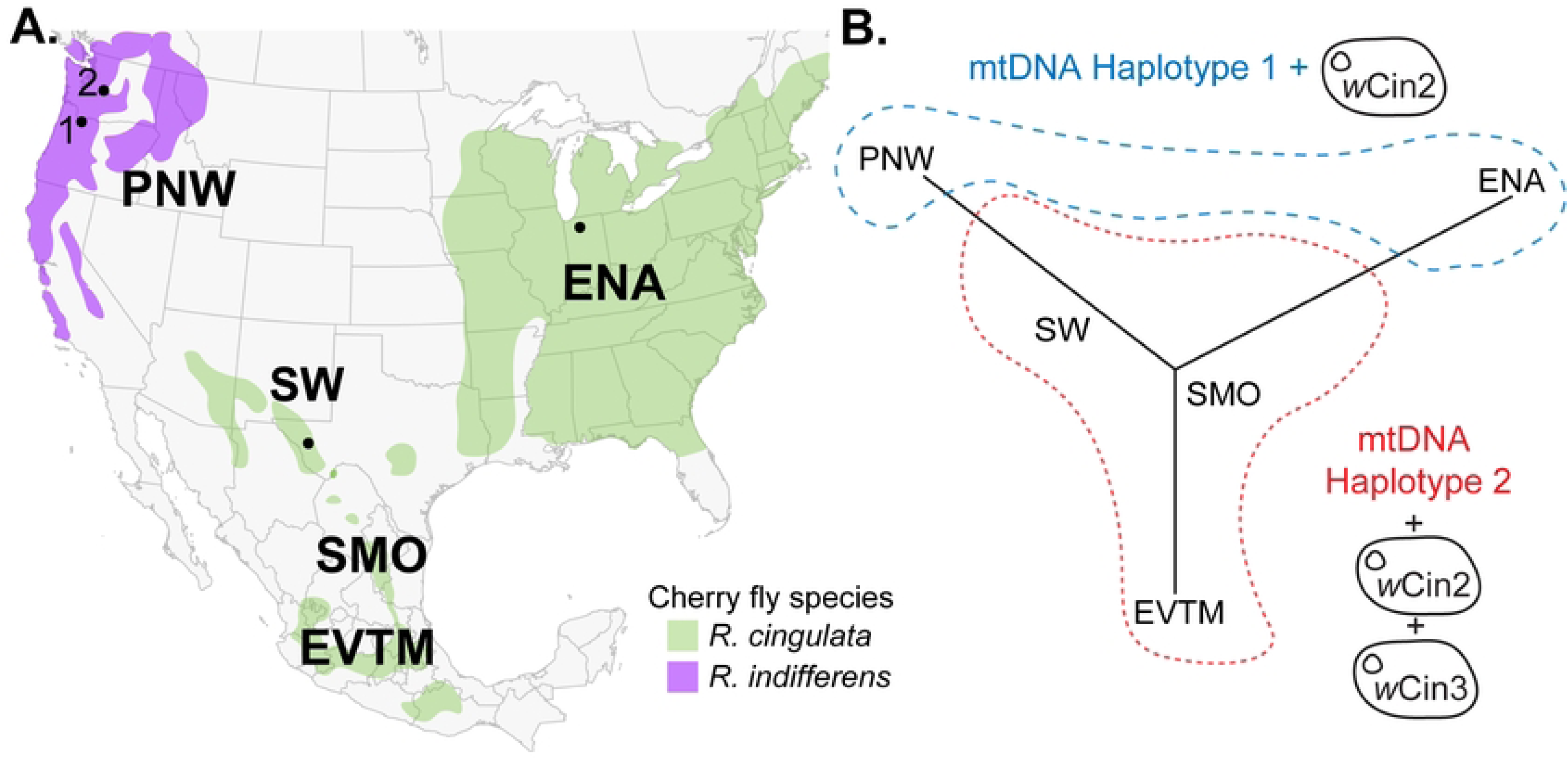
Cherry fly populations and their genotypes. (A) Map of *R. cingulata* and *R. indifferens* populations from Eastern N. America (ENA), the Southwest (SW), Sierra Madre Oriental Mountains (SMO), the Eje Volcánico Transversal de Mexicano (EVTM), and two populations from the Pacific Northwest (PNW) (S1 Table). (B) A neighbor-joining tree of *R. cingulata* and *R. indifferens* from Doellman et al. (2020) [50] superimposed with mtDNA haplotypes (haplotype 1 in blue and haplotype 2 in red) and known associated *Wolbachia* strains.

The disjunct geographic distribution of mtDNA haplotypes in cherry flies coincides with a difference in *Wolbachia* strain composition. All cherry fly individuals across N. America are infected by the same group A *Wolbachia* strain, *w*Cin2 [18,51,52]. However, Multi Locus Sequence Typing (MLST) indicates that *R. cingulata* flies in the SW, SMO, and EVTM are also coinfected by a second, highly diverged group B strain, *w*Cin3 [18]. Crosses between doubly infected SW to singly infected PNW and ENA cherry flies show a pattern consistent with *w*Cin3 inducing unidirectional CI [18,48]. It has therefore been hypothesized that a cherry fly population in Mexico became coinfected with the CI-inducing *w*Cin3 strain from a yet to be identified source 100,000 to 157,000 years ago, resulting in its association with the derived mtDNA haplotype. Subsequently, *w*Cin3 spread northward carrying the disjunct haplotype but was halted before it could reach the PNW or ENA due to climate change in the Holocene [18]. Hitchhiking of a mitochondrial haplotype with a spreading *Wolbachia* strain is not uncommon and is found in several other insect systems [53,54].

Given their recent Holocene separation across N. America and the more distant mtDNA divergence associated with the acquisition and sweep of *w*Cin3, the *R. cingulata* species group presents a model system for studying *Wolbachia* genome dynamics across different evolutionary timescales. Here, we leverage this natural history to investigate the dynamics of genome evolution for the universal *w*Cin2 *Wolbachia* strain that diverged with its cherry fly hosts across three timeframes: (1) Recent: between two populations of *R. indifferens* infesting bitter cherry, isolated by distance in the PNW by < 1,000 years; (2) Intermediate: between populations of *R. indifferens* in the PNW and *R. cingulata* infesting black cherry in the ENA, estimated to have an early Holocene divergence; and (3) Distant: between the *R. cingulata* populations in the SW versus *R. indifferens* in the PNW and *R. cingulata* in the ENA, where strains have been separated by ≥ 100,000 years. Using Nanopore sequencing, we assembled complete *Wolbachia* genomes from these host fly populations and assessed them for differences in gene sequence content, *cif* genes, MGEs, and genome structure to characterize *Wolbachia* genome evolution over varying temporal scales. While earlier MLST analyses suggested that *w*Cin2 strains from the ENA and PNW were identical, with *w*Cin2 from the SW differing by only a single SNP [18], our whole genome approach found that *w*Cin2 strains are far more diverse than previously thought. The assembled *w*Cin2 genomes revealed differences in sequence identity, gene content, *cif* genes, MGEs, synteny, and structural rearrangements, with levels of differentiation increasing with time since separation. Crucially, our findings indicate that *Wolbachia* genomes, influenced by MGEs, can evolve rapidly and impact *cif* genes and subsequent host diversification via CI.

## Results

### Long-read sequencing generated six closed *Wolbachia* genomes

*Wolbachia* genomes were assembled with 25X to 50X Oxford Nanopore Technologies (ONT) (Oxford, UK) long reads generated from high molecular weight (HMW) DNA extracted from individual flies from the PNW1, PNW2, and SW populations (Fig 1 and S1 Table). Additionally, we included the completed 1.53 Mb *w*Cin2-ENA genome (GCF_017604245.1 [52]) in our analyses. All *Wolbachia* genomes assembled into single circular contigs with Benchmarking Universal Single-Copy Orthologs (BUSCO) scores greater than 98.70% based on the Rickettsiales dataset (Fig 2 and S2 Table). For the PNW1 population, a single 1.53 Mb *w*Cin2-PNW1 strain was assembled. For the PNW2 population, two coinfecting *Wolbachia* strains were assembled: a 1.53 Mb *w*Cin2-PNW2 stain and a new, previously undescribed, 1.24 Mb *Wolbachia* strain, designated *w*Ind (Fig 2). For the SW population, two coinfecting *Wolbachia* strains were also assembled: a 1.56 Mb *w*Cin2-SW strain and a 1.52 Mb *w*Cin3 strain. *In silico* annotations show that MGEs, including transposases, recombinases, and resolvases, make up a sizable portion of these assembled genomes, from 10.74% to 12.39% (S3 Table). We identified eight prophage regions in *w*Cin2-ENA, *w*Cin2-PNW1, and *w*Cin2-PNW2, but only three prophage regions in *w*Cin2-SW (Fig 2). Additionally, the *w*Cin3 strain coinfecting flies in the SW contained eight prophage regions, while the *w*Ind strain coinfecting PNW2 flies contained two prophage regions (Fig 2).

**Fig 2.**
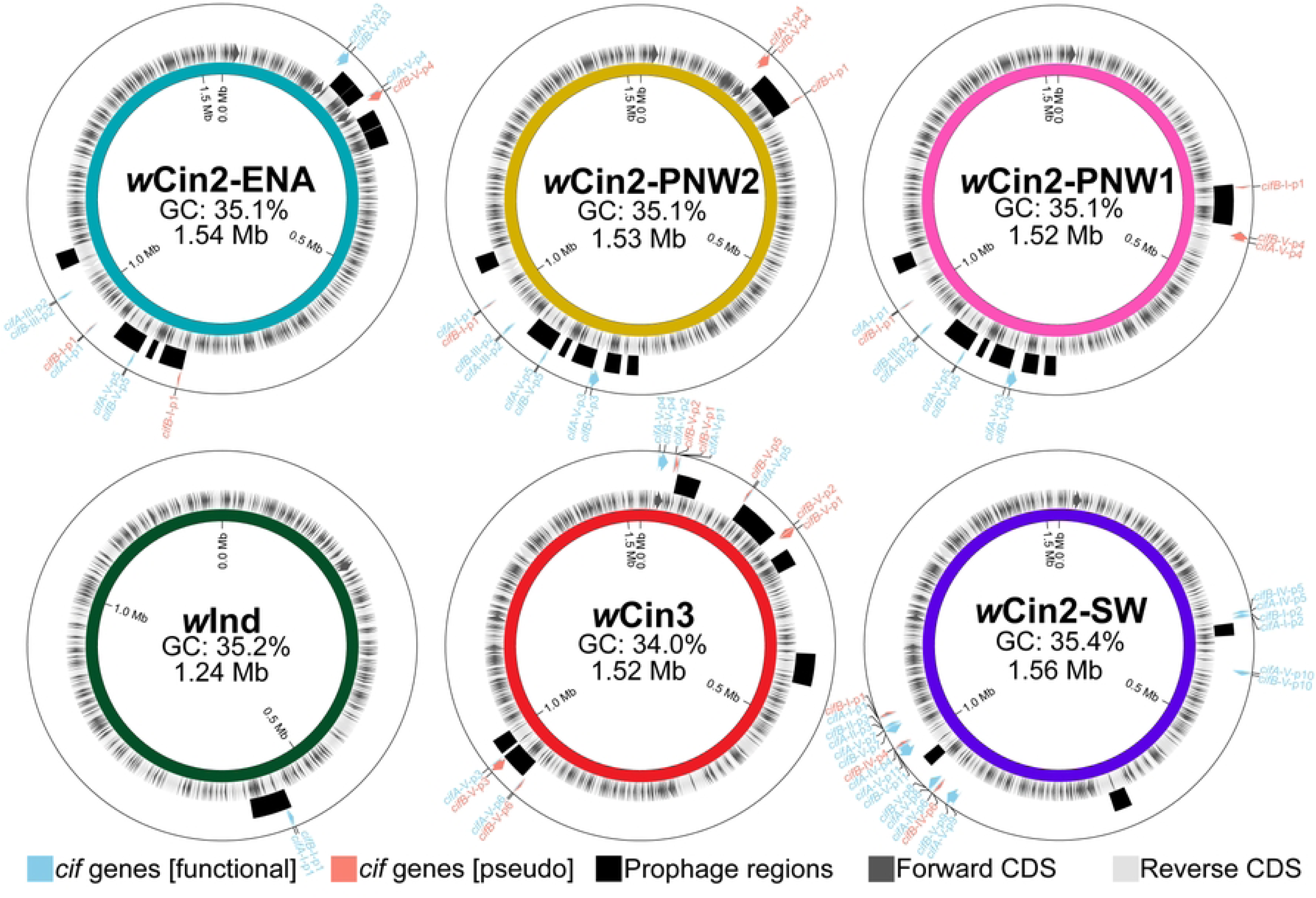
Six closed *Wolbachia* genomes were assembled from four cherry fly populations. The inner ring represents forward and reverse coding sequences. The middle ring highlights predicted prophage regions. The outermost ring depicts functional and pseudogenized *cif* genes. See S2 Table for additional assembly statistics and for NCBI accession numbers.

### 33 pairs of *cif* genes were identified

To catalogue the *cif* gene repertoire of the assembled cherry fly *Wolbachia* genomes, we used protein BLAST (BLASTp) searches using reference *cif* homologues (S4 Table). These searches identified 33 *cif* gene pairs representing all five major *cif* gene types in the assembled cherry fly *Wolbachia* genomes (Fig 2 and S5 Table). Many functional and several pseudogenized *cif* genes were found, the latter indicated by the presence of missense frameshift mutations and stop codons. In general, *cifB* genes were more likely to be pseudogenized than *cifA* genes, agreeing that selection acts to maintain CI rescue functions but not CI induction [29,41,55]. Out of the 14 possible pseudogenized *cif* genes identified, only two were *cifA* genes compared to 12 *cifB* genes.

Strains *w*Cin2-ENA, *w*Cin2-PNW1, and *w*Cin2-PNW2 shared five pairs of identical *cif* genes, comprising one pseudogenized pair of *cif_[T1]_*, one pair of *cif_[T3]_*, and three pairs of *cif_[T5]_* genes, with one of the three pairs of *cif_[T5]_* genes predicted to be pseudogenized (Fig 2, S5 Table, S1 Fig, and S2 Fig). The pair of *cif_[T1]_* genes had high identities (99%) to the *Drosophila cif_wMel[T1]_* gene but likely do not induce CI as the *cifB_[T1]_*genes were truncated and split into two fragments, with the second fragment containing an internal stop codon. Furthermore, *cifB_[T1]_* in *w*Cin2-ENA was upstream of *cifA_[T1]_*, preventing joint transcription of *cifA* and *cifB*. In *w*Cin2-PNW1 and *w*Cin2-PNW2, the *cifB_[T1]_*was translocated several hundred kilobases away from its paired *cifA_[T1]._* The pair of *cif_[T3]_* genes had high identities (99% and 97%) to *cif_wNol[T3]_,* are likely functional, and are in an inversion in the *w*Cin2-PNW1 and *w*Cin2-PNW2 strains compared to *w*Cin2-ENA (Fig 2 and S5 Table). Two pairs of *cif_[T5]_* genes were also predicted to be functional with identities closest to *cif_wTri-2[T5]_* and *cif_wStri-1[T5]_*, respectively. Finally, one pair of *cif_[T5]_* genes with closest identities to *cif_wStri-2[T5]_* were pseudogenized with stop codons found in both the *cifA* and *cifB* genes. Only the *cifA_wStri-2[T5]_* gene from *w*Cin2-ENA appeared functional.

Strain *w*Cin2-SW had 11 pairs of *cif* genes, possessing: two *cif_[T1]_* pairs, one *cif_[T2]_* pair, three *cif_[T4]_* pairs, and five *cif_[T5]_* pairs (Fig 2, S5 Table, S1 Fig, and S2 Fig). The first *cif_[T1]_* pair had high identities (99%) to *cif_wMel[T1]_*, but a stop codon pseudogenized the *cifB_[T1]_* gene. The second pair *cif_[T1]_* pair was derived with low identities (65% and 61%) to *cif_wMel[T1]_* but appeared functional. The *cif_[T2]_* pair were also derived with low identities (58% and 56%) to *cif_wRi[T2]_* and was predicted to be functional. The three *cif_[T4]_* pairs had identities ranging from 97% to 52% to *cif_wAlbB[T4]_.* However, only one of the three *cif_[T4]_* pairs was functional, with the other two pairs containing internal stop codons in the *cifB_[T4]_* gene. All five *cif_[T5]_* pairs had identities ranging from 84% to 56% to *cif_wTri-2[T5]_*, *cif_Stri-1[T5]_*, *cif_wStri-2[T5]_*, or *cif_wStri-1[T5]_* and appeared functional (Fig 2 and S5 Table).

Strain *w*Ind infecting *R. indifferens* in the PNW2 population had a pair of syntenic *cif_[T1]_* genes displaying 61% sequence identity to *cifA_wMel[T1]_* and 60% identity to *cifB_wMel[T1]_*, respectively (Fig 2, S5 Table, S1 Fig, and S2 Fig). Strain *w*Cin3 infecting *R. cingulata* in the SW had six pairs of *cif_[T5]_*genes, with identities ranging from 93% to 43% identity to *cif_wTri-2[T5]_*, *cif_Stri-1[T5]_*, *cif_wStri-2[T5]_*or *cif_wStri-1[T5]_.* While all the *cifA_[T5]_* genes were functional, only one of six were paired with a functional *cifB* gene. The remaining five *cifB* genes were pseudogenized containing stop codons and/or frameshift mutations (Fig 2 and S5 Table).

### *w*Cin3 and *w*Ind are highly diverged from *w*Cin2

To place the newly assembled *w*Cin3 and *w*Ind *Wolbachia* strains in a larger phylogenetic context, we constructed a RAxML tree based on 208 single-copy orthologous loci shared among all our sequenced strains and other representative A and B supergroup *Wolbachia* strains (Fig 3). Strain *w*Cin3, coinfecting *R. cingulata* flies from the SW and Mexico, was inferred to be within the *Wolbachia* B supergroup. The closest related strain to *w*Cin3 was *w*AlbB, a *Wolbachia* strain that infects *Aedes albopictus*. Strain *w*Ind was genetically diverged from the other *w*Cin2 strains and placed at the base of the *Wolbachia* A supergroup. The closest relatives to *w*Ind are the *Wolbachia* strain *w*Ano62 that infects ants [56] and *w*Orie and *w*Neo, that infect *Drosophila* species [29]. All the *w*Cin2 strains clustered with the *w*Mel clade within the A supergroup.

**Figure 3.**
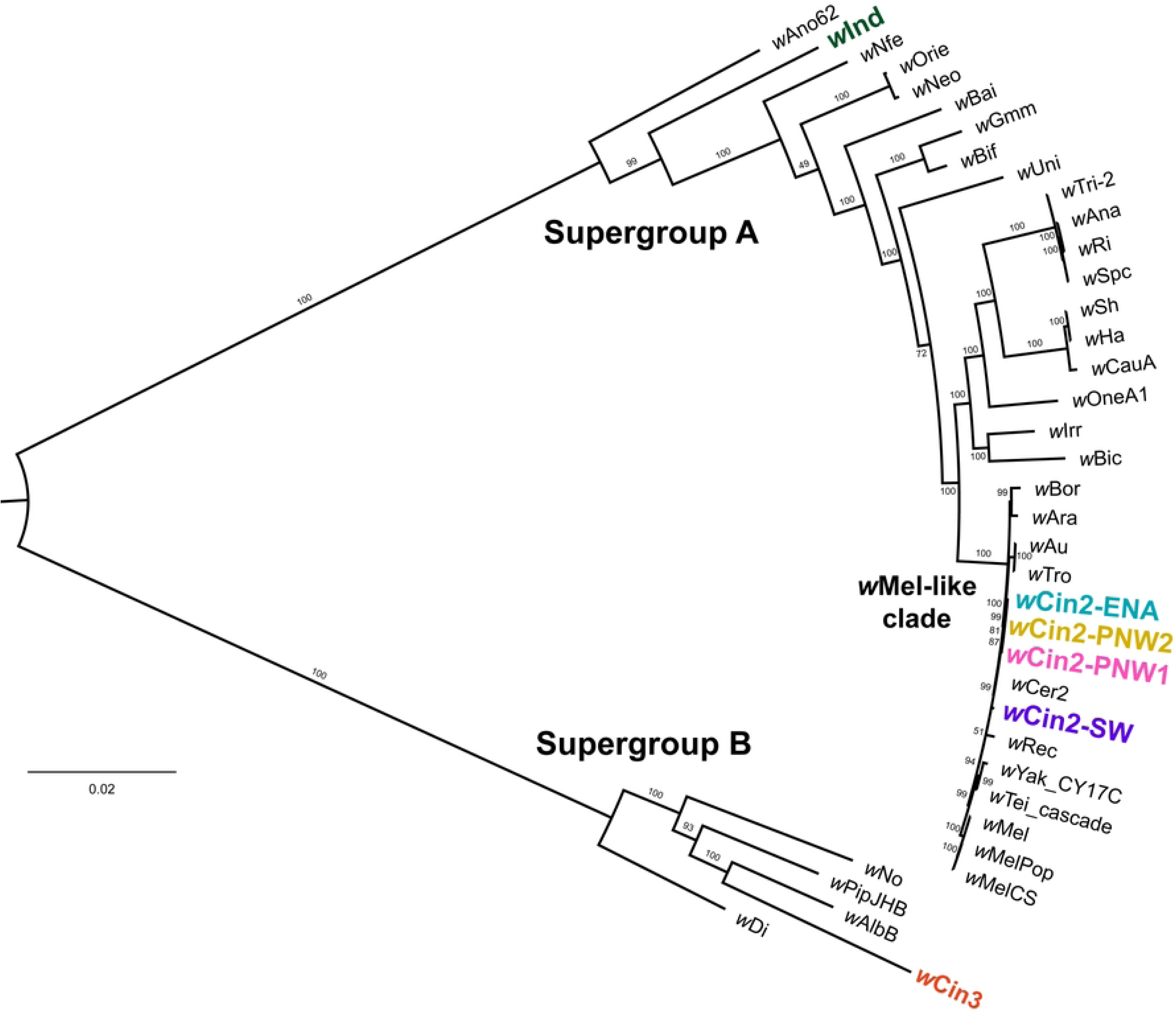
*Wolbachia* RAxML tree shows *w*Cin3 and *w*Ind are diverged from *w*Cin2. The A and B supergroup *Wolbachia* RAxML tree was derived from a set of 208 single-copy orthologous shared genes. Bootstrap support values are given for nodes based on 1,000 replicates. The cherry fly infecting *Wolbachia* strains are highlighted with different colors. Metadata for the A and B supergroup reference genomes are found in the S6 Table.

### *w*Cin2-SW is diverged from other *w*Cin2 strains

To resolve the evolutionary relationships among *w*Cin2 strains, a RAxML tree was constructed that was limited to A supergroup *w*Mel-like *Wolbachia* and cherry fly *w*Cin2 strains using a shared set of 975 single-copy orthologous genes (Fig 4A). Here, all *w*Cin2 strains in *R. cingulata* and *R. indifferens* clustered and were sister to *w*Mel strains from *Drosophila* (Fig 4A). In addition, strain *w*Cin2-SW was genetically distinct from the *w*Cin2-ENA, *w*Cin2-PNW1, and *w*Cin2-PNW2 strains (Fig 4A). Mean sequence similarity between the *w*Cin2-ENA, *w*Cin2-PNW1, and *w*Cin2-PNW2 strains was very high (>99.85%), resulting in a polytomy in the RAxML tree (Fig 4B). Pairwise comparisons showed the highest sequence similarity between *w*Cin2-ENA and *w*Cin2-PNW2 strains (99.92%) and between *w*Cin2-PNW1 and *w*Cin2-PNW2 strains (99.91%). Sequence similarity was lower for all pairwise comparisons involving the ENA and PNW strains to *w*Cin2-SW (>98.21%). The RAxML tree for *w*Cin2 therefore concurred with the disjunct pattern displayed by cherry fly mtDNA haplotypes and differed from the clinal pattern shown by host microsatellites, with a genetic break seen for *w*Cin2 between the ENA and PNW versus the SW [50].

**Figure 4.**
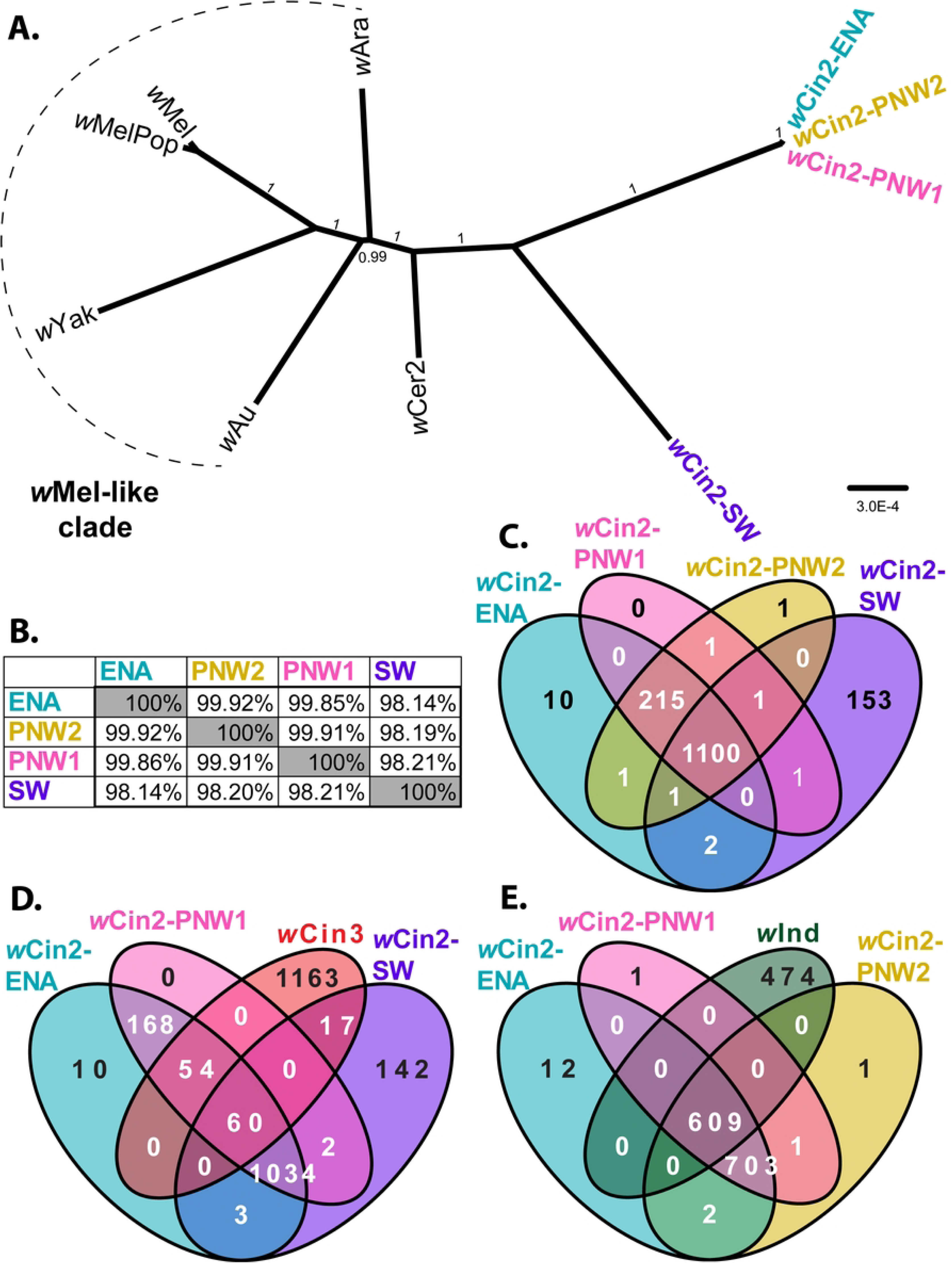
*w*Cin2-SW is diverged from the other *w*Cin2 strains. (A) RAxML tree for the clade of *w*Mel-like *Wolbachia* strains based on 975 single-copy shared orthologous genes with 1,000 bootstraps. The *w*Cin2 strains are highlighted with different colors. (B) Pairwise percent identity for the four *w*Cin2 genomes that infect cherry flies. (C) Venn diagram detailing the number of unique genes for each *w*Cin2 strain taken from the 1,486 genes in the *w*Cin2 pangenome. (D) Venn diagram showing 17 unique genes shared between the co-infecting *Wolbachia* strains *w*Cin3 and *w*Cin2-SW and 54 unique genes shared between *w*Cin3 and wCn2-ENA/PNW1, not found in *w*Cin2-SW. (E) Venn diagram showing 0 unique genes shared between *w*Ind and *w*Cin2-PNW2 as well as the other *w*Cin2 strains.

Analysis of a set of 1,486 loci shared amongst *w*Cin2 strains indicated that the genome of *w*Cin2-SW not only differed in sequence content but also differed significantly in gene content from the other strains (Fig 4C). Overall, 1,100 genes were shared among the four *w*Cin2 strains. There were no unique genes identified in the genome of *w*Cin2-PNW1, one unique gene was found in *w*Cin2-PNW2, and 10 unique genes were detected in *w*Cin2-ENA. In contrast, 153 unique genes were identified in *w*Cin2-SW. Furthermore, *w*Cin2 from the ENA, PNW1, and PNW2 shared 215 unique genes not found in the *w*Cin2-SW strain. Most of the unique genes from each strain had annotations related to MGEs (S3 Fig). For example, of the 10 genes unique to *w*Cin2-ENA, five were *in silico* annotated and four of these were classified as MGEs (S7 Table). Of the 153 unique genes from *w*Cin2-SW, 40 of the 64 annotated genes were classified as MGEs (S8 Table). Finally, of the 215 unique genes shared among *w*Cin2-ENA, *w*Cin2-PNW1 and *w*Cin2-PNW2, 59 of the 101 annotated genes were classified as MGEs (S9 Table).

Comparisons of shared and unique genes between the *w*Cin2 strains and *w*Cin3 yielded surprising results that suggest the possibility of past genetic exchange between the coinfecting strains. A total of 17 genes were shared only between *w*Cin3 and *w*Cin2-SW (Fig 4D and S10 Table). Notably, *cifA_wStri-2[T5]_* and a *cifB_wStri-2[T5]_*genes were identified as being unique to both strains, clustering together in their respective *cifA* and *cifB* RAxML trees (Figs S1 and S2). We also identified 54 genes shared only between *w*Cin3, *w*Cin2-ENA, and *w*Cin2-PNW1, not found in *w*Cin2-SW (Fig 4D). Again, most of these shared genes were classified as MGEs and related to prophages (S11 Table). In contrast, we found no unique gene shared between coinfecting strains *w*Ind and *w*Cin2-PNW2 and no unique gene shared between *w*Ind and *w*Cin2-PNW1 and *w*Cin2-ENA (Fig 4E).

### Structural variation differentiates closely related *w*Cin2 strains

Despite having near-identical gene content and sequence similarity, we detected unique structural variation between *w*Cin2-ENA, *w*Cin2-PNW1, and *w*Cin2-PNW2 (Fig 5). Two major genomic translocations and a large inversion differentiated the *w*Cin2-ENA from the PNW strains. The two PNW *w*Cin2 stains, *w*Cin2-PNW1 and *w*Cin2-PNW2, also differed from each other by a large 344 kb inversion (Fig 5). To determine the genes associated with these structural variants, we manually searched for annotated genes contained within the inversions and translocations. Each inversion and translocation contained MGEs, including prophages and transposases, as well as *cif* genes. One of the translocations between *w*Cin2-ENA and the PNW *w*Cin2 contained a complete prophage and a pair of *cif_[T5]_* genes. The second, smaller translocation between *w*Cin2-ENA and PNW *w*Cin2 contained a *cifA_[T1]_* gene. The inversion between *w*Cin2-ENA and PNW *w*Cin2 strains contained a pair of *cif_[T5]_* genes and an unpaired *cifB_[T1]_*. The large inversion between *w*Cin2-PNW1 and *w*Cin2-PNW2 contained a pair of *cif_[T5],_* genes, an unpaired *cifA_[T1]_*, and a complete prophage. Whole genome alignments were also performed between the *w*Cin2-ENA, *w*Cin2-PNW2, and *w*Cin2-PNW1 strains versus *w*Cin2-SW, resulting in the identification of over 51 rearrangements between them (S4 Fig).

**Figure 5.**
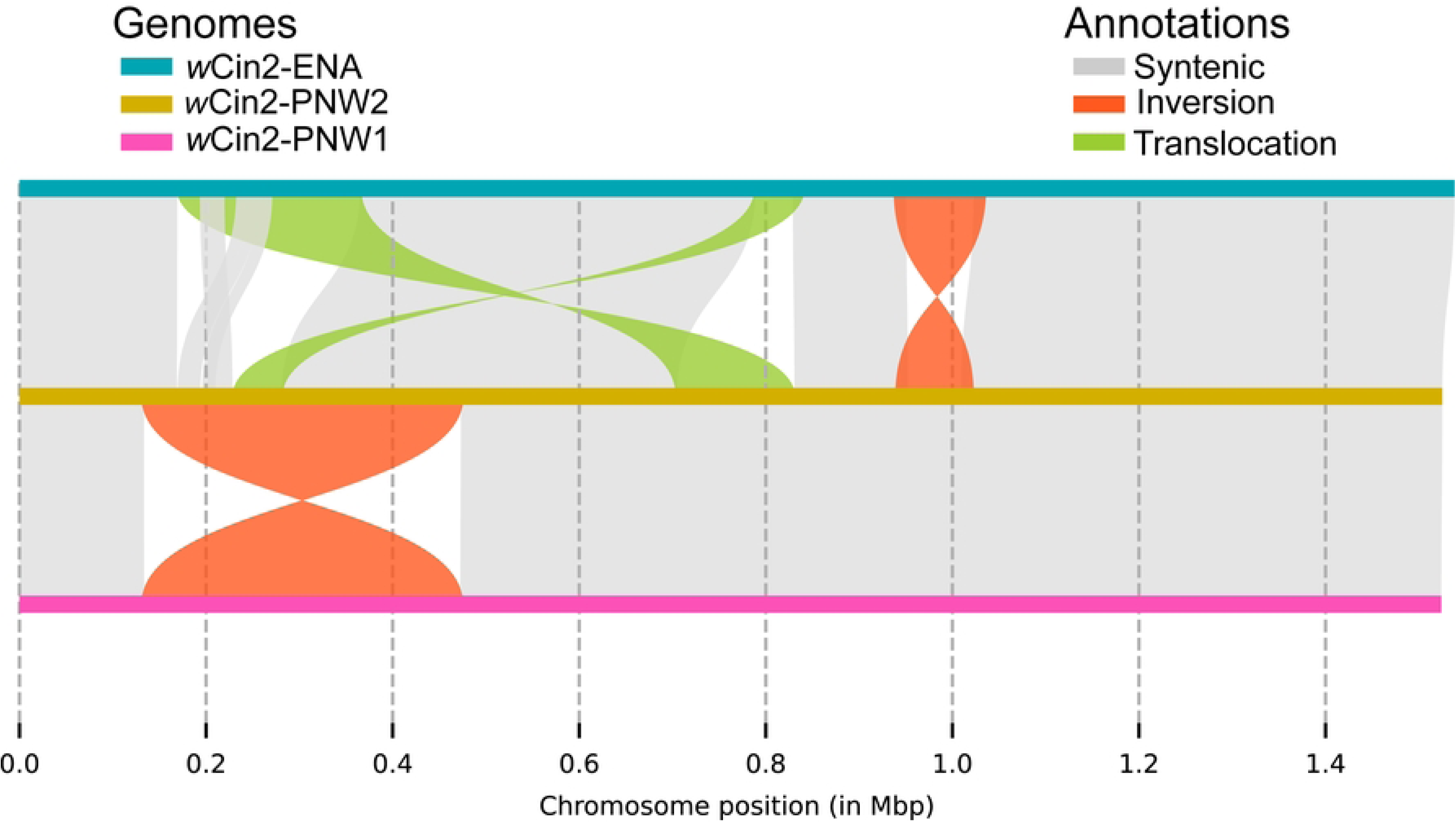
Synteny plots detail structural differences between the closely related *w*Cin2 strains. Whole genome alignments found that two translocations and an inversion differentiate the *w*Cin2-ENA and the *w*Cin2 strains from the PNW. The *w*Cin2-PNW1 and *w*Cin2-PNW2 strains are differentiated by a large 344 kb inversion.

### Within population differentiation of *Wolbachia* strains

To investigate potential intra-strain variability, we conducted nanopore sequencing on two to three additional samples from each population and performed structural variant scans. Our analysis revealed that the major translocations and inversions identified above, which distinguish *w*Cin2 strains, were also present in the additional sequenced individuals and thus appear fixed at the population level. Besides these fixed inversions and translocations, we identified several indels segregating among the *w*Cin2 samples within each population. Most of the detected indels were less than 50 bp, although indels greater than 50 bp were present in nine out of ten samples (S12 Table). Consequently, while each sequenced *w*Cin2 sample shared population-specific structural variants, each also exhibited a unique set of indels.

## Discussion

In this study, we characterized the evolution of the *Wolbachia* strain, *w*Cin2, present in all cherry-infesting populations of *Rhagoletis* in the *R. cingulata* sibling species group. We utilized nanopore sequencing to assemble closed, circular, genomes for *Wolbachia* from flies from four different populations across N. America that diverged from less than 1,000 to more than 100,000 years ago. This approach enabled comparative genomic analysis of *Wolbachia* sequence divergence, gene content, synteny, and structure across evolutionary timescales ranging from the recent to the more distant past. While earlier MLST analyses suggested that *w*Cin2 strains from the ENA and PNW were identical, with *w*Cin2 from the SW differing by only a single SNP [18], our whole genome approach revealed that *w*Cin2 strains are far more diverse than previously thought. The assembled *w*Cin2 genomes exhibited substantial differences in sequence identity, gene content, *cif* genes, MGEs, and structural rearrangements, with levels of differentiation increasing with time since separation.

Most importantly, our findings show that *Wolbachia* genome structure can rapidly evolve. On a recent timescale of less than 1,000 years, genomic sequences for *w*Cin2 strains from the two cherry fly populations in the Pacific Northwest (PNW1 & PNW2) were near-identical. However, a notable structural variation was observed where *w*Cin2-PNW1 and *w*Cin2-PNW2 differ by a 344 kb inversion, flipping two prophage regions and a *cif* gene pair. At an intermediate timescale of several thousand years separating cherry flies from the PNW and ENA, genome sequences were also near-identical for the *w*Cin2 strains. Nevertheless, small differences in gene content were observed between the *w*Cin2 from the ENA and PNW, including 10 unique genes in *w*Cin2-ENA and a pseudogenized *cifA_[T5]_*in the PNW, the latter of which appears functional in *w*Cin2-ENA. Most significantly, two translocations and a small inversion further distinguished *w*Cin2 strains from the ENA and PNW populations.

With additional nanopore sequencing of two to three additional individuals per population, we confirmed that the major structural variants noted between the *w*Cin2 genomes appear to be fixed at the population level, at least in the samples we assessed. These results are consistent with other recent studies suggesting that population structure may be common in some *Wolbachia* strains [36,57]. In addition to the fixed structural rearrangements distinguishing *w*Cin2 strains between populations, we identified polymorphic deletions and insertions in *w*Cin2 genomes within populations, most of which were under 1000 bp. Indeed, within a fly population, every sequenced host fly exhibited a unique set of indels in its associated *w*Cin2 genome. Intra-population structural variants have recently been identified as mutants in lab populations of *w*Mel and *w*Pip [43–45], but our results show that intra-population structural variants are likely prevalent in wild populations too. Our findings highlight the dynamic evolution of *Wolbachia*, and rather than a fixed strain, *Wolbachia* invasions could be made of up of a group of highly similar variants. Population-level sampling will be important to fully understand *Wolbachia* strain diversity in the future.

The implications of rapid structural evolution of *Wolbachia* genomes over short to modest time periods are even more pronounced when considering strains that diverged over much longer time scales. In this regard, our results align with previous studies, demonstrating that once *Wolbachia* strains exhibit measurable levels of sequence divergence, their gene content and genome structure are typically already highly differentiated [29,37,39,42,58,59]. The *w*Cin2 strain from the SW, which became associated with a unique mtDNA haplotype over 100,000 years ago, differs by an estimated 51 translocations and inversions from wCin2 genomes in the PNW and ENA, with very few regions showing synteny. Gene content also rapidly diversified in *w*Cin2-SW, with the strain containing 153 unique genes, a reduction in the number of prophage regions from eight to three, and the presence of 11 pairs of *cif* genes — five more pairs than any other *w*Cin2 strain. Notably, by our estimates the 11 pairs of *cif* genes in *w*Cin2-SW are the highest number identified in a single *Wolbachia* strain to date [29,37]. These findings in cherry flies thus echo the broader pattern exhibited in other systems, where over longer evolutionary time periods, the gain and loss of *cif* genes decouples them from *Wolbachia* and host divergence patterns, often leading to the creation of new CI phenotypes [38,41,42].

Our findings provide further evidence that MGEs constitute a substantial portion of the *Wolbachia* genome and significantly contribute to variation in gene content and structure among closely related strains [36,52,60–65]. In our assembled *Wolbachia* genomes, more than 10% of genes were annotated as transposases, and each genome contained between two and eight predicted active prophages. Additionally, MGEs can be disruptive force and lead to the pseudogenization of *Wolbachia* genes [66,67]. In our study, all major structural rearrangements detected in *w*Cin2 involved MGEs, either containing prophages or transposons. We also identified several instances of pseudogenized *cif* genes resulting from MGE associated translocations and inversions, including a *cifB* gene removed from its *cif*A partner.

Furthermore, MGEs often move between different *Wolbachia* strains and can facilitate the transfer of nearby genes, most notably *cif* genes [29,38,68]. We found evidence for gene transfer events between *w*Cin3 and *w*Cin2-SW, with 17 unique genes shared between them, most of which were prophage related. The high number of *cif* genes found in *w*Cin2-SW could partly be attributed to transfers from the coinfecting *w*Cin3 strain, or perhaps another unknown, previous coinfecting strain. Specifically, we discovered that both the *w*Cin2-SW and the coinfecting strain *w*Cin3 possess a *cif_[wDacT5]_* gene pair not found in any of the other *w*Cin2 strains from the ENA or PNW. Finally, we observed that most *cif* gene pairs in *w*Cin2 were flanked by transposases, likely aiding in their intra- and inter-genomic transfers [38]. These findings underscore the significant role MGEs play in shaping *Wolbachia* genomes, driving genetic diversity and adaptability through rapid structural evolution and gene transfer events.

Previous work suggested that the unidirectional reduction in egg hatch rate observed in cherry fly crosses between SW males to ENA and PNW females was due to CI induced by *w*Cin3, a supergroup B strain coinfecting SW flies along with *w*Cin2 [18,48]. However, our genomic results suggest a possible alternative scenario, where *w*Cin2-SW may also play a role in causing CI. The genome of *w*Cin3 currently contains six pairs of *cif_[T5]_* genes, with only a single *cif_[T5]_* gene pair appearing functional. While *cif_[T5]_* homologs have been shown to induce CI [29], they are present in all the *w*Cin2 strains, including those infecting cherry flies from the ENA and PNW. Given that related *cif* homologs can generally rescue CI [13,63], this implies that *w*Cin3 alone could not be responsible for causing the observed CI in crosses between male SW cherry flies to PNW and ENA females [18]. It remains possible, however, that CI in cherry flies is caused by strains infected with *cif_[T5]_* genes from different clades. Martinez *et al*. (2020) showed that the *cif_[T5]_* clades are as diverse as the *cif_[T1]_, cif_[T2]_, cif_[T3]_, and cif_[T4]_* clades, and perhaps like these clades, *cif_[T5]_* clades may be able to induce and rescue their own specific type of CI. Notably, we found that strain *w*Cin2-SW and the coinfecting strain *w*Cin3 both possess a *cif_[wDacT5]_* gene pair not found in the other *w*Cin2 strains. This suggests that *cif_[wDacT5]_*may have been acquired in *w*Cin2-SW through inter-strain transfer from *w*Cin3, and together both strains could now be contributing to CI. Alternatively, *w*Cin2-SW alone could be responsible for the observed CI. Strain *w*Cin2-SW harbors 11 pairs of *cif* genes, eight of which are putatively functional. Two types of *cif* genes present in *w*Cin2-SW, *cif_[T2]_* and *cif_[T4]_*, are absent in other *w*Cin2 strains from the PNW1, PNW2, and ENA. One pair of *cif_[T4]_* and one pair of *cif_[T2]_* homologs are particularly significant because *in silico* annotations predict them to be fully functional, with no evidence of pseudogenization.

Genome comparisons also revealed that *w*Cin2-ENA, *w*Cin2-PNW1, and *w*Cin2-PNW2 all share a unique functional pair of *cif_[T3]_* homologs not found in *w*Cin2-SW or *w*Cin3. It is possible that CI may also be induced by *cif_[T3]_* genes from *w*Cin2-ENA, *w*Cin2-PNW1, and *w*Cin2-PNW2. Empirical evidence, however, does not support this scenario, as crosses between ENA and PNW males to SW females show no evidence for reduced egg hatch [18]. This discrepancy suggests that CI induction by the *cif_[T3]_* homologs might be rescued by one of the other *cif* homologs in *w*Cin2-SW or *w*Cin3, or is due to some other factor such as *cif* transcript levels [31] or other *Wolbachia cif-*associated genes [63].

In summary, our examination of the *Wolbachia* strain *w*Cin2, which infests cherry fly populations ranging in divergence times from the recent to the more distant past, has provided significant insights into the rapid and complex evolutionary dynamics of *Wolbachia* genomes. Our findings demonstrate that *Wolbachia* genomes, driven by MGEs, can evolve swiftly, leading to substantial differences in sequence identity, gene content (including *cif* genes), and most prominently structural rearrangements. This work highlights the value of using long-read sequencing to assemble completed genomes needed to unravel the intricate and often subtle genomic changes that occur within *Wolbachia* strains and populations. Future research should focus on further characterizing the functional roles of the identified *cif* genes in *w*Cin2 and *w*Cin3 to better understand how CI is induced by SW *R. cingulata* males, potentially using techniques such as antisense RNA to selectively silence specific *cif* genes [69]. By continuing to compare *Wolbachia* strains and *cif* gene dynamics across different evolutionary timescales and host populations, we can deepen our understanding of the evolutionary processes that govern endosymbiont-host relationships, with broader implications for adaptation and speciation.

## Methods

### Fly collection and ONT sequencing

Specimens of *R. cingulata* and *R. indifferens* were collected from 2018 to 2021 as larvae feeding in infested *P. serotina* fruit from the eastern USA (ENA), southwest USA (SW), and from *P. emarginata* fruit from two sites in the Pacific Northwest USA (PNW): Hood River, Oregon (PNW1) and Cle Elum, Washington (PNW2) (Fig 1A; S1 Table). Larvae were reared to adulthood following standard *Rhagoletis* husbandry techniques, with a 6–8 month overwintering treatment at 4 °C [18,48,70,71]. Newly eclosed adults were isolated by sex and matured for a minimum of seven days in cages with food (1-part autolyzed yeast to 3-parts honey) and water. Mature females were frozen and stored at -80 °C until sequencing.

High molecular weight DNA was extracted from individual flies following Wolfe et al. [52]. ONT LSK109 ligation sequencing libraries were prepared according to manufacturer’s instructions from induvial flies and sequenced on R9.4.1 flow cells using a MinION Mk1B device. For cherry fly populations from PNW1, PNW2, and SW (the *w*Cin2 from ENA has already been published: GCF_017604245.1 [52]), we first sequenced one female fly at high coverage (25X-50X *Wolbachia* reads) to generate closed *Wolbachia* genomes. After this, two additional flies were sequenced from each population at a low coverage (∼10X *Wolbachia* reads) to search for any missed structural variants at the population level (S1 Table). Finally, 150 bp paired-end Illumina libraries using the same DNA from the high coverage individuals were prepared by the Notre Dame Genomics Core and sequenced on a HiSeq X Ten platform at BGI Genomics to polish final *Wolbachia* assemblies.

### *Wolbachia* genome assembly

ONT signal data were basecalled using Guppy v6.01 with the SUP model and gently quality filtered (>Q8 and >500 bp) with nanoq [72]. For the high coverage sequencing runs used for genome assembly, reads were mapped with minimap2 [73] to the *w*Cin2 reference genome (GCF_017604245.1) and assembled using Flye v2.9.1 [74]. Assemblies were first polished using Racon v1.4.13 [75] followed by medaka 1.5.0. Short reads, quality filtered (>Q20 and >80 bp) with fastp [76], were used to further polish ONT assemblies with four iterations of Pilon [77]. Circlator [78] synchronized the start site for all the completed assemblies to *dnaA*. Assemblies were evaluated for completeness using BUSCO v4 with the *Rickettsiales* dataset [79].

### Annotation of *Wolbachia* genomes and *cif* genes

Completed *Wolbachia* assemblies were first annotated with Prokka v1.14.6 [80] using Pfam, PGAP, and HAMAP protein databases [81–83]. Phage regions were annotated using PHASTEST [84]. Finally, *cif* genes were identified using representative *cifA* and *cifB* genes from each of the five phylogenetic types as query sequences for BLASTp searches [28,29] (S4 Table). Positive BLASTp hits were accepted if E-values were close to 0, identity was greater than 40%, and the positive hit was greater than 40% of the queries length. Putative *cif* candidates were evaluated for pseudogenization using Artemis [85] to manually check for insertions, premature stop codons, or frameshift mutations and to evaluate if the *cifA* and *cifB* genes were in a complete module, with *cifB* immediately downstream from *cifA*. The above gene, phage, and *cif* gene annotations were visualized with pyCirclize [86].

### *cif* gene phylogenies

Predicted *cifA* and *cifB* genes identified from the above BLASTp searches and the BLASTp *cif* references (S4 Table) were aligned with MAFFT [87]. Both the derived *cifA* and *cifB* pseudogenes and the derived *w*Bor reference *cif* genes were excluded to improve phylogenetic signal. The *cifA* and *cifB* amino acid alignments were used to generate *cifA* and *cifB* RAxML-NG v1.1 [88] trees using the LG+G8+F substitution model with 25 random and 25 parsimony-based starting trees and 1,000 bootstrap replicates.

### *Wolbachia* phylogenetic analyses

Phylogenetic analysis was performed at two scales, involving (1) both A and B supergroup *Wolbachia* strains; and (2) considering only *w*Mel-like A supergroup strains. Panaroo v1.3 [89] found sets of orthologous single-copy core genes using the annotations produced by Prokka from the genomes assembled above and a group of reference genomes (S6 Table). The data set for A and B supergroups together included 208 single-copy orthologous genes (387,932 bp) and the dataset for A group *w*Mel-like *Wolbachia* alone included 975 orthologous single-copy genes (924,342 bp). For each dataset, gene sequences were concatenated and aligned using MAFFT [87] and RAxML-NG v1.1 [88] constructed phylogenetic trees using the GTR+G substitution model, with 50 random and 50 parsimony-based starting trees, and with 1,000 bootstrap replicates.

### *Wolbachia* strain *w*Cin2 genome comparisons

Shared and unique genes for *w*Cin2 strains were determined from a set of single-copy orthologous genes. Using the previously generated Prokka annotations, Panaroo generated a pangenome for all the 1,486 protein coding genes for all four *w*Cin2 strains. Using this dataset, R was used to determine the unique and shared genes between each strain [90]. Pairwise whole genome comparisons were performed for all *w*Cin2 genomes using MUMmer4 [91] to estimate average percent gene identity. MUMmer4 was also used to generate whole-genome alignments between strains that had the highest identity to one another to detect structural rearrangements (PNW1 to PNW2 and ENA to PNW2). Synteny and any rearrangements detected between these alignments were annotated with SyRI [92] and plotted with Plotsr [93]. SyRI annotations were validated against results produced by Mauve [94]. We were unable to use SyRI for comparisons to the *w*Cin2-SW genome because of its higher divergence, and instead used Mauve [94].

To verify whether the identified structural variations were representative of each *w*Cin2 population, we checked for additional structural variants from two to three additional samples sequenced at low coverage. We used CuteSV2, shown to have high accuracy at estimating structural variants (SVs) with low coverage data [95], to scan the ONT reads for any additional synteny not captured in our assembled reference genomes. For each population, ONT reads were aligned to their reference genome using minimap2 and SVs were called using cuteSV2 with settings optimized for ONT data with a minimum of five reads needed to call a breakpoint.

## Acknowledgements

We would like to thank Jaqueline Lopez and the Note Dame Genomics and Bioinformatics core facility for the preparation of Illumina libraries and DNA quality control, as well as Mike Pfrender and Stuart Jones for insightful comments on early drafts. We also thank Robert B Goughnour, Glen Ray Hood, Meredith M. Doellman, and Cheyenne Tait for fly collection and/or husbandry for samples sequenced in the study.

## Supporting information

**S1 Table. Collection metadata for sequenced flies.**

Populations are shown in Fig. 1. Flies were sampled from 2018-2021 from the ENA, the SW, and two populations in the PNW – one from Hood River, Oregon (PNW1) and one from Cle Elum, Washington State (PNW2). Samples sequenced at high coverage (∼30-50X *Wolbachia* reads) were used for genome assembly and samples sequenced at low coverage (∼10X *Wolbachia* reads) were used for structural variant scans. GCF_017604245.1 is the ENA-HC sample and the *w*Cin2 reference genome.

**S2 Table. Genome assembly stats for cherry fly associated *Wolbachia* strains.**

BUSCO scores from the Rickettsiales dataset. Assembly stats are for all assembled *Wolbachia* genomes, including the reference *w*Cin2 genome (GCF_017604245.1) from the ENA.

**S3 Table. MGEs are highly prevalent in the assembled *Wolbachia* genomes.**

Counts of the number of *in silico* Bakta annotations that are classified of MGEs compared to the total number of coding genes in the genome

**S4 Table. Representative *cif* genes for each type (I-V) were used for reference sequences for BLASTp searches of the assembled *Wolbachia* genomes.**

Listed are the *Wolbachia* strains associated with each *cif* gene type and associated NCBI gene ID.

**S5 Table. BLASTp hit table of putative *cif* homologous for all assembled *Wolbachia* strains.**

Representative *cif* genes from S1 Table were used as reference sequences. Hit start and Hit end denote the start and stop coordinates of the *cif* genes in assembled genomes, direction describes the coding direction of the *cif* genes, and e-val quantifies the significance of the BLASTp hits. Gene identities quantify similarity between the reference *cif* gene and the *cif* BLASTp hits.

**S6 Table. Reference genomes used for phylogenetic analyses.**

The table lists the *Wolbachia* strain, its supergroup, the RefSeq assembly ID, genome size, number contigs, and associated references.

**S7 Table. Annotations for the 10 unique genes for *w*Cin2-ENA.**

All genes for *w*Cin2-ENA were annotated with Prokka using the Pfam, PGAP, and HAMAP protein databases. Then using *R,* the genes unique to *w*Cin2-ENA were sorted.

**S8 Table. Annotations for the 153 unique genes for *w*Cin2-SW.**

All genes for *w*Cin2-SW were annotated with Prokka using the Pfam, PGAP, and HAMAP protein databases. Then using *R,* the genes unique to *w*Cin2-SW were sorted.

**S9 Table. Annotations for the 215 unique genes shared between *w*Cin2-ENA, *w*Cin2-PNW1, and *w*Cin2-PNW2.**

All genes within the shared gene set were annotated with Prokka using the Pfam, PGAP, and HAMAP protein databases. Then using *R,* the genes shared between *w*Cin2-ENA, *w*Cin2-PNW1, and *w*Cin2-PNW2 were sorted.

**AS10 Table. Annotations for the 17 unique genes shared between *w*Cin2-SW and *w*Cin3.**

All genes within the shared gene set were annotated with Prokka using the Pfam, PGAP, and HAMAP protein databases. Most annotated genes are MGEs.

**S11 Table. Annotations for the 54 unique genes shared between *w*Cin3. *w*Cin2-ENA, and *w*Cin2-PNW1.**

All genes within the shared gene set were annotated with Prokka using the Pfam, PGAP, and HAMAP protein databases. Most annotated genes are MGEs.

**S12 Table. Several small SVs are found at the population level for the *w*Cin2 strains.**

Low coverage SV scans using CuteSV against the assembled wCin2 genomes identified several SVs less than 1000 bp and four SVs between 1000 and 10,000 bp

**S1 Fig. The cherry fly *cifA* gene RAxML phylogeny highlights *cifA* gene diversity.**

The RAxML phylogeny was made using a MAFFT amino acid alignment with 1000 bootstraps. Derived pseudogenes were removed before constructing the phylogeny. Strain *w*Cin2-SW has the most *cifA* genes with 11 copies, *w*Cin3 has six copies, *w*Cin2-ENA has five copies, and the *w*Cin2 strains from the PNW have four functional copies. See S5 Table for more detailed *cif* gene information.

**S2 Fig. The cherry fly *cifB* gene RAxML phylogeny highlights *cifB* gene diversity.**

The RAxML phylogeny was made using a MAFFT amino acid alignment with 1000 bootstraps. Derived pseudogenes were removed before constructing the phylogeny, those that remain are denoted with **. Strain *w*Cin2-SW has the most *cifB* functional copies with eight, followed by the ENA and PNW strains with two each, and further followed still by *w*Cin3 and *w*Ind with one copy. See S5 Table for more detailed *cif* gene information.

**S3 Fig. Annotated genes show enrichment for MGE genes in *w*Cin2 strains.**

Most annotated genes unique to either the *w*Cin2-ENA, *w*Cin2-PNW1/PNW2/ENA, or *w*Cin2-SW strains are MGEs.

**S4 Fig. Synteny comparisons between the *w*Cin2 strains show *w*Cin2-SW is highly diverged.**

Structural genome rearrangements were identified with Mauve. Strain *w*Cin2-SW has over 51 structural rearrangements compared to *w*Cin2-ENA. The PNW *w*Cin2 strains only have four and three rearrangements compared to *w*Cin2-ENA.

## References

1. Weinert LA, Araujo-Jnr EV, Ahmed MZ, Welch JJ. The incidence of bacterial endosymbionts in terrestrial arthropods. Proc Biol Sci. 2015;282: 20150249. doi:10.1098/rspb.2015.0249

2. Taylor MJ, Bandi C, Hoerauf A. *Wolbachia* bacterial endosymbionts of filarial nematodes. Adv Parasitol. 2005;60: 245–284. doi:10.1016/S0065-308X(05)60004-8

3. Baldo L, Ayoub NA, Hayashi CY, Russell JA, Stahlhut JK, Werren JH. Insight into the routes of *Wolbachia* invasion: high levels of horizontal transfer in the spider genus *Agelenopsis* revealed by *Wolbachia* strain and mitochondrial DNA diversity. Mol Ecol. 2008;17: 557–569. doi:10.1111/j.1365-294X.2007.03608.x

4. Sanaei E, Charlat S, Engelstädter J. *Wolbachia* host shifts: routes, mechanisms, constraints and evolutionary consequences. Biol Rev Camb Philos Soc. 2021;96: 433–453. 10.1111/brv.12663

5. Dedeine F, Vavre F, Fleury F, Loppin B, Hochberg ME, Boulétreau M. Removing symbiotic *Wolbachia* bacteria specifically inhibits oogenesis in a parasitic wasp. Proc Natl Acad Sci U S A. 2001;98: 6247–6252. doi:10.1073/pnas.101304298

6. Hosokawa T, Koga R, Kikuchi Y, Meng X-Y, Fukatsu T. *Wolbachia* as a bacteriocyte-associated nutritional mutualist. Proc Natl Acad Sci U S A. 2010;107: 769–774. doi:10.1073/pnas.0911476107

7. Newton ILG, Rice DW. The jekyll and hyde symbiont: could *Wolbachia* be a nutritional mutualist? J Bacteriol. 2020;202: e00589–19. doi:10.1128/JB.00589-19

8. Hedges LM, Brownlie JC, O’Neill SL, Johnson KN. *Wolbachia* and virus protection in insects. Science. 2008;322: 702–702. doi:10.1126/science.1162418

9. Teixeira L, Ferreira Á, Ashburner M. The bacterial symbiont *Wolbachia* induces resistance to RNA viral infections in *Drosophila melanogaster*. PLoS Biol. 2008;6: e1000002. doi:10.1371/journal.pbio.1000002

10. Cogni R, Ding SD, Pimentel AC, Day JP, Jiggins FM. *Wolbachia* reduces virus infection in a natural population of *Drosophila*. Commun Biol. 2021;4: 1–7. doi:10.1038/s42003-021-02838-z

11. Kaur R, Shropshire JD, Cross KL, Leigh B, Mansueto AJ, Stewart V, et al. Living in the endosymbiotic world of *Wolbachia*: A centennial review. Cell Host Microbe. 2021;29: 879–893. doi:10.1016/j.chom.2021.03.006

12. Correa CC, Ballard JWO. *Wolbachia* associations with insects: winning or losing against a master manipulator. Front Ecol Evol. 2016;3: 153. doi:10.3389/fevo.2015.00153

13. Shropshire JD, Leigh B, Bordenstein SR. Symbiont-mediated cytoplasmic incompatibility: What have we learned in 50 years? eLife. 2020;9: e61989. doi:10.7554/eLife.61989

14. Telschow A, Hammerstein P, Werren JH. The effect of *Wolbachia* on genetic divergence between populations: models with two-way migration. Am Nat. 2002;160: S54–S66. doi:10.1086/342153

15. Flor M, Hammerstein P, Telschow A. *Wolbachia*-induced unidirectional cytoplasmic incompatibility and the stability of infection polymorphism in parapatric host populations. J Evol Biol. 2007;20: 696–706. 10.1111/j.1420-9101.2006.01252.x

16. Brucker RM, Bordenstein SR. Speciation by symbiosis. Trends Ecol Evol. 2012;27: 443–451. doi:10.1016/j.tree.2012.03.011

17. Cruz MA, Magalhães S, Sucena É, Zélé F. *Wolbachia* and host intrinsic reproductive barriers contribute additively to postmating isolation in spider mites. Evolution. 2021;75: 2085–2101. doi:10.1111/evo.14286

18. Bruzzese DJ, Schuler H, Wolfe TM, Glover MM, Mastroni JV, Doellman MM, et al. Testing the potential contribution of *Wolbachia* to speciation when cytoplasmic incompatibility becomes associated with host-related reproductive isolation. Mol Ecol. 2022;31: 2935–2950. doi:10.1111/mec.16157

19. Telschow A, Hammerstein P, Werren JH. The effect of *Wolbachia* versus genetic incompatibilities on reinforcement and speciation. Evolution. 2005;59: 1607–1619. doi:10.1111/j.0014-3820.2005.tb01812.x

20. Jaenike J, Dyer KA, Cornish C, Minhas MS. Asymmetrical reinforcement and *Wolbachia* infection in *Drosophila*. PLoS Biol. 2006;4: e325. doi:10.1371/journal.pbio.0040325

21. Beckmann JF, Ronau JA, Hochstrasser M. A *Wolbachia* deubiquitylating enzyme induces cytoplasmic incompatibility. Nat Microbiol. 2017;2: 17007. doi:10.1038/nmicrobiol.2017.7

22. Chen H, Ronau JA, Beckmann JF, Hochstrasser M. A *Wolbachia* nuclease and its binding partner provide a distinct mechanism for cytoplasmic incompatibility. Proc Natl Acad Sci U S A. 2019;116: 22314–22321. doi:10.1073/pnas.1914571116

23. LePage DP, Metcalf JA, Bordenstein SR, On J, Perlmutter JI, Shropshire JD, et al. Prophage WO genes recapitulate and enhance *Wolbachia* -induced cytoplasmic incompatibility. Nature. 2017;543: 243–247. doi:10.1038/nature21391

24. McNamara CJ, Ant TH, Harvey-Samuel T, White-Cooper H, Martinez J, Alphey L, et al. Transgenic expression of *cif* genes from *Wolbachia* strain *w*AlbB recapitulates cytoplasmic incompatibility in *Aedes aegypti*. Nat Commun. 2024;15: 869. doi:10.1038/s41467-024-45238-7

25. Shropshire JD, Bordenstein SR. Two-By-One model of cytoplasmic incompatibility: Synthetic recapitulation by transgenic expression of *cifA* and *cifB* in *Drosophila*. PLoS Genet. 2019;15: e1008221. doi:10.1371/journal.pgen.1008221

26. Shropshire JD, On J, Layton EM, Zhou H, Bordenstein SR. One prophage WO gene rescues cytoplasmic incompatibility in *Drosophila melanogaster*. Proc Natl Acad Sci U S A. 2018;115: 4987– 4991. doi:10.1073/pnas.1800650115

27. Adams KL, Abernathy DG, Willett BC, Selland EK, Itoe MA, Catteruccia F. *Wolbachia cifB* induces cytoplasmic incompatibility in the malaria mosquito vector. Nat Microbiol. 2021;6: 1575–1582. doi:10.1038/s41564-021-00998-6

28. Lindsey ARI, Rice DW, Bordenstein SR, Brooks AW, Bordenstein SR, Newton ILG. Evolutionary genetics of cytoplasmic incompatibility genes *cifA* and *cifB* in Prophage WO of *Wolbachia*. Genome Biol Evol. 2018;10: 434–451. doi:10.1093/gbe/evy012

29. Martinez J, Klasson L, Welch JJ, Jiggins FM. Life and death of selfish genes: comparative genomics reveals the dynamic evolution of cytoplasmic incompatibility. Mol Biol Evol. 2021;38: 2–15. doi:10.1093/molbev/msaa209

30. Reynolds KT, Thomson LJ, Hoffmann AA. The effects of host age, host nuclear background and temperature on phenotypic effects of the virulent *Wolbachia* strain popcorn in *Drosophila melanogaster*. Genetics. 2003;164: 1027–1034. doi:10.1093/genetics/164.3.1027

31. Shropshire JD, Hamant E, Conner WR, Cooper BS. *cifB*-transcript levels largely explain cytoplasmic incompatibility variation across divergent *Wolbachia*. PNAS Nexus. 2022;1: pgac099. doi:10.1093/pnasnexus/pgac099

32. Wybouw N, Mortier F, Bonte D. Interacting host modifier systems control *Wolbachia*-induced cytoplasmic incompatibility in a haplodiploid mite. Evol Lett. 2022;6: 255–265. doi:10.1002/evl3.282

33. Klasson L, Westberg J, Sapountzis P, Näslund K, Lutnaes Y, Darby AC, et al. The mosaic genome structure of the *Wolbachia w*Ri strain infecting *Drosophila simulans*. Proc Natl Acad Sci U S A. 2009;106: 5725–5730. doi:10.1073/pnas.0810753106

34. Tanaka K, Furukawa S, Nikoh N, Sasaki T, Fukatsu T. Complete WO phage sequences reveal their dynamic evolutionary trajectories and putative functional elements required for integration into the *Wolbachia* genome. Appl Environ Microbiol. 2009;75: 5676–5686. doi:10.1128/AEM.01172-09

35. Kent BN, Bordenstein SR. Phage WO of *Wolbachia*: lambda of the endosymbiont world. Trends Microbiol. 2010;18: 173–181. doi:10.1016/j.tim.2009.12.011

36. Hill T, Unckless RL, Perlmutter JI. Positive selection and horizontal gene transfer in the genome of a male-killing *Wolbachia*. Mol Biol Evol. 2022;39: msab303. doi:10.1093/molbev/msab303

37. Martinez J, Ant TH, Murdochy SM, Tong L, Filipe A da S, Sinkins SP. Genome sequencing and comparative analysis of *Wolbachia* strain *w*AlbA reveals *Wolbachia*-associated plasmids are common. PLoS Genet. 2022;18: e1010406. doi:10.1371/journal.pgen.1010406

38. Cooper BS, Vanderpool D, Conner WR, Matute DR, Turelli M. *Wolbachia* acquisition by *Drosophila yakuba*-clade hosts and transfer of incompatibility loci between distantly related *Wolbachia*. Genetics. 2019;212: 1399–1419. doi:10.1534/genetics.119.302349

39. Scholz M, Albanese D, Tuohy K, Donati C, Segata N, Rota-Stabelli O. Large scale genome reconstructions illuminate *Wolbachia* evolution. Nat Commun. 2020;11: 5235. doi:10.1038/s41467-020-19016-0

40. Collier LS, Largaespada DA. Transposable elements and the dynamic somatic genome. Genome Biol. 2007;8: S5. doi:10.1186/gb-2007-8-s1-s5

41. Turelli M, Cooper BS, Richardson KM, Ginsberg PS, Peckenpaugh B, Antelope CX, et al. Rapid global spread of *w*Ri-like *Wolbachia* across multiple *Drosophila*. Curr Biol. 2018;28: 963–971.e8. doi:10.1016/j.cub.2018.02.015

42. Meany MK, Conner WR, Richter SV, Bailey JA, Turelli M, Cooper BS. Loss of cytoplasmic incompatibility and minimal fecundity effects explain relatively low *Wolbachia* frequencies in *Drosophila mauritiana*. Evolution. 2019;73: 1278–1295. doi:10.1111/evo.13745

43. Duarte EH, Carvalho A, López-Madrigal S, Costa J, Teixeira L. Forward genetics in *Wolbachia*: Regulation of *Wolbachia* proliferation by the amplification and deletion of an addictive genomic island. PLoS Genet. 2021;17: e1009612. doi:10.1371/journal.pgen.1009612

44. Chrostek E, Teixeira L. Mutualism Breakdown by Amplification of *Wolbachia* Genes. PLoS Biol. 2015;13: e1002065. doi:10.1371/journal.pbio.1002065

45. Namias A, Ngaku A, Makoundou P, Unal S, Sicard M, Weill M. Intra-lineage microevolution of *Wolbachia* leads to the emergence of new cytoplasmic incompatibility patterns. PLoS Biol. 2024;22: e3002493. doi:10.1371/journal.pbio.3002493

46. Bush GL. The taxonomy, cytology, and evolution of the genus *Rhagoletis* in North America (Diptera, Tephritidae). Bull Mus Comp Zool. 1966;134: 431–562.

47. Bush GL. Sympatric host race formation and speciation in frugivorous flies of the genus *Rhagoletis* (Diptera, Tephritidae). Evolution. 1969;23: 237–251. doi:10.2307/2406788

48. Tadeo E, Feder JL, Egan SP, Schuler H, Aluja M, Rull J. Divergence and evolution of reproductive barriers among three allopatric populations of *Rhagoletis cingulata* across eastern North America and Mexico. Entomol Exp Appl. 2015;156: 301–311. doi:10.1111/eea.12331

49. Doellman MM, Schuler H, Jean GS, Hood GR, Egan SP, Powell THQ, et al. Geographic and ecological dimensions of host plant-associated genetic differentiation and speciation in the *Rhagoletis cingulata* (Diptera: Tephritidae) sibling species group. Insects. 2019;10: 275. doi:10.3390/insects10090275

50. Doellman MM, Jean GS, Egan SP, Powell THQ, Hood GR, Schuler H, et al. Evidence for spatial clines and mixed geographic modes of speciation for North American cherry-infesting *Rhagoletis* (Diptera: Tephritidae) flies. Ecol Evol. 2020;10: 12727–12744. 10.1002/ece3.6667

51. Schuler H, Bertheau C, Egan SP, Feder JL, Riegler M, Schlick-Steiner BC, et al. Evidence for a recent horizontal transmission and spatial spread of *Wolbachia* from endemic *Rhagoletis cerasi* (Diptera: Tephritidae) to invasive *Rhagoletis cingulata* in Europe. Mol Ecol. 2013;22: 4101–4111. doi:10.1111/mec.12362

52. Wolfe TM, Bruzzese DJ, Klasson L, Corretto E, Lečić S, Stauffer C, et al. Comparative genome sequencing reveals insights into the dynamics of *Wolbachia* in native and invasive cherry fruit flies. Mol Ecol. 2021;30: 6259–6272. doi:10.1111/mec.15923

53. Hurst GDD, Jiggins FM. Problems with mitochondrial DNA as a marker in population, phylogeographic and phylogenetic studies: the effects of inherited symbionts. Proc R Soc Lond B Biol Sci. 2005;272: 1525–1534. doi:10.1098/rspb.2005.3056

54. Schuler H, Köppler K, Daxböck-Horvath S, Rasool B, Krumböck S, Schwarz D, et al. The hitchhiker’s guide to Europe: the infection dynamics of an ongoing *Wolbachia* invasion and mitochondrial selective sweep in *Rhagoletis cerasi*. Mol Ecol. 2016;25: 1595–1609. doi:10.1111/mec.13571

55. Beckmann JF, Vaerenberghe KV, Akwa DE, Cooper BS. A single mutation weakens symbiont-induced reproductive manipulation through reductions in deubiquitylation efficiency. Proc Natl Acad Sci U S A. 2021;118. doi:10.1073/pnas.2113271118

56. Lee C-C, Lin C-Y, Tseng S-P, Matsuura K, Yang C-CS. Ongoing coevolution of *Wolbachia* and a widespread invasive ant, *Anoplolepis gracilipes*. Microorganisms. 2020;8: 1569. doi:10.3390/microorganisms8101569

57. Kaur R, Siozios S, Miller WJ, Rota-Stabelli O. Insertion sequence polymorphism and genomic rearrangements uncover hidden *Wolbachia* diversity in *Drosophila suzukii* and *D. subpulchrella*. Sci Rep. 2017;7: 14815. doi:10.1038/s41598-017-13808-z

58. Vancaester E, Blaxter M. Phylogenomic analysis of *Wolbachia* genomes from the Darwin Tree of Life biodiversity genomics project. PLoS Biol. 2023;21: e3001972. doi:10.1371/journal.pbio.3001972

59. Gao S, Ren Y-S, Su C-Y, Zhu D-H. High levels of multiple phage WO infections and its evolutionary dynamics associated with *Wolbachia*-infected butterflies. Front Microbiol. 2022;13: 865227. doi:10.3389/fmicb.2022.865227

60. Ishmael N, Hotopp JCD, Ioannidis P, Biber S, Sakamoto J, Siozios S, et al. Extensive genomic diversity of closely related *Wolbachia* strains. Microbiology. 2009;155: 2211–2222. doi:10.1099/mic.0.027581-0

61. Ellegaard KM, Klasson L, Näslund K, Bourtzis K, Andersson SGE. Comparative genomics of *Wolbachia* and the bacterial species concept. PLoS Genet. 2013;9: e1003381. doi:10.1371/journal.pgen.1003381

62. Gerth M, Bleidorn C. Comparative genomics provides a timeframe for *Wolbachia* evolution and exposes a recent biotin synthesis operon transfer. Nat Microbiol. 2016;2: 1–7. doi:10.1038/nmicrobiol.2016.241

63. Baião GC, Janice J, Galinou M, Klasson L. Comparative genomics reveals factors associated with phenotypic expression of *Wolbachia*. Genome Biol Evol. 2021;13: evab111. doi:10.1093/gbe/evab111

64. Wu M, Sun LV, Vamathevan J, Riegler M, Deboy R, Brownlie JC, et al. Phylogenomics of the reproductive parasite *Wolbachia pipientis w*Mel: a streamlined genome overrun by mobile genetic elements. PLoS Biol. 2004;2: e69. doi:10.1371/journal.pbio.0020069

65. Duplouy A, Iturbe-Ormaetxe I, Beatson SA, Szubert JM, Brownlie JC, McMeniman CJ, et al. Draft genome sequence of the male-killing *Wolbachia* strain *w*Bol1 reveals recent horizontal gene transfers from diverse sources. BMC Genomics. 2013;14: 20. doi:10.1186/1471-2164-14-20

66. Sanogo YO, Dobson SL, Bordenstein SR, Novak RJ. Disruption of the *Wolbachia* surface protein gene wspB by a transposable element in mosquitoes of the *Culex pipiens* complex (Diptera, Culicidae). Insect Mol Biol. 2007;16: 143–154. doi:10.1111/j.1365-2583.2006.00707.x

67. Queffelec J, Postma A, Allison JD, Slippers B. Remnants of horizontal transfers of *Wolbachia* genes in a *Wolbachia*-free woodwasp. BMC Ecol Evo. 2022;22: 36. doi:10.1186/s12862-022-01995-x

68. Wang GH, Sun BF, Xiong TL, Wang YK, Murfin KE, Xiao JH, et al. Bacteriophage WO can mediate horizontal gene transfer in endosymbiotic *Wolbachia* genomes. Front Microbiol. 2016;7: 1867. doi:10.3389/fmicb.2016.01867

69. Hussain M, Zhang G, Leitner M, Hedges LM, Asgari S. *Wolbachia* RNase HI contributes to virus blocking in the mosquito *Aedes aegypti*. iScience. 2022;26: 105836. doi:10.1016/j.isci.2022.105836

70. Feder JL, Chilcote CA, Bush GL. Inheritance and linkage relationships of allozymes in the apple maggot fly. J Hered. 1989;80: 277–283. doi:10.1093/oxfordjournals.jhered.a110854

71. Yee WL, Goughnour RB, Hood GR, Forbes AA, Feder JL. Chilling and host plant/site-associated eclosion times of Western cherry fruit fly (Diptera: Tephritidae) and a host-specific parasitoid. Environ Entomol. 2015;44: 1029–1042. doi:10.1093/ee/nvv097

72. Steinig E, Coin L. Nanoq: ultra-fast quality control for nanopore reads. J Open Source Softw. 2022;7: 2991. doi:10.21105/joss.02991

73. Li H. Minimap2: pairwise alignment for nucleotide sequences. Bioinformatics. 2018;34: 3094– 3100. doi:10.1093/bioinformatics/bty191

74. Kolmogorov M, Yuan J, Lin Y, Pevzner PA. Assembly of long, error-prone reads using repeat graphs. Nat Biotechnol. 2019;37: 540–546. doi:10.1038/s41587-019-0072-8

75. Vaser R, Sovic I, Nagarajan N, Sikic M. Fast and accurate de novo genome assembly from long uncorrected reads. Genome Res. 2017; gr.214270.116. doi:10.1101/gr.214270.116

76. Chen S, Zhou Y, Chen Y, Gu J. fastp: an ultra-fast all-in-one FASTQ preprocessor. Bioinformatics. 2018;34: i884–i890. doi:10.1093/bioinformatics/bty560

77. Walker BJ, Abeel T, Shea T, Priest M, Abouelliel A, Sakthikumar S, et al. Pilon: An integrated tool for comprehensive microbial variant detection and genome assembly improvement. PLoS One. 2014;9: e112963. doi:10.1371/journal.pone.0112963

78. Hunt M, Silva ND, Otto TD, Parkhill J, Keane JA, Harris SR. Circlator: automated circularization of genome assemblies using long sequencing reads. Genome Biol. 2015;16: 294. doi:10.1186/s13059-015-0849-0

79. Manni M, Berkeley MR, Seppey M, Simão FA, Zdobnov EM. BUSCO update: novel and streamlined workflows along with broader and deeper phylogenetic coverage for ccoring of Eukaryotic, Prokaryotic, and viral genomes. Mol Biol Evol. 2021;38: 4647–4654. doi:10.1093/molbev/msab199

80. Seemann T. Prokka: rapid prokaryotic genome annotation. Bioinformatics. 2014;30: 2068–2069. doi:10.1093/bioinformatics/btu153

81. Pedruzzi I, Rivoire C, Auchincloss AH, Coudert E, Keller G, de Castro E, et al. HAMAP in 2015: updates to the protein family classification and annotation system. Nucleic Acids Res. 2015;43: D1064–D1070. doi:10.1093/nar/gku1002

82. Tatusova T, DiCuccio M, Badretdin A, Chetvernin V, Nawrocki EP, Zaslavsky L, et al. NCBI prokaryotic genome annotation pipeline. Nucleic Acids Res. 2016;44: 6614–6624. doi:10.1093/nar/gkw569

83. Mistry J, Chuguransky S, Williams L, Qureshi M, Salazar GA, Sonnhammer ELL, et al. Pfam: The protein families database in 2021. Nucleic Acids Res. 2021;49: D412–D419. doi:10.1093/nar/gkaa913

84. Wishart DS, Han S, Saha S, Oler E, Peters H, Grant JR, et al. PHASTEST: faster than PHASTER, better than PHAST. Nucleic Acids Res. 2023;51: W443–W450. doi:10.1093/nar/gkad382

85. Carver T, Harris SR, Berriman M, Parkhill J, McQuillan JA. Artemis: an integrated platform for visualization and analysis of high-throughput sequence-based experimental data. Bioinformatics. 2012;28: 464–469. doi:10.1093/bioinformatics/btr703

86. Shimoyama Y. pyCirclize: Circular visualization in Python. 2022. Available: https://github.com/moshi4/pyCirclize

87. Katoh K, Misawa K, Kuma K, Miyata T. MAFFT: a novel method for rapid multiple sequence alignment based on fast Fourier transform. Nucleic Acids Res. 2002;30: 3059–3066. doi:10.1093/nar/gkf436

88. Kozlov AM, Darriba D, Flouri T, Morel B, Stamatakis A. RAxML-NG: a fast, scalable and user-friendly tool for maximum likelihood phylogenetic inference. Bioinformatics. 2019;35: 4453–4455. doi:10.1093/bioinformatics/btz305

89. Tonkin-Hill G, MacAlasdair N, Ruis C, Weimann A, Horesh G, Lees JA, et al. Producing polished prokaryotic pangenomes with the Panaroo pipeline. Genome Biol. 2020;21: 180. doi:10.1186/s13059-020-02090-4

90. R Core Team. R: a language and environment for statistical computing. Vienna, Austria: R Foundation for Statistical Computing; 2019. Available: https://www.R-project.org/

91. Marçais G, Delcher AL, Phillippy AM, Coston R, Salzberg SL, Zimin A. MUMmer4: A fast and versatile genome alignment system. PLoS Comput Biol. 2018;14: e1005944. doi:10.1371/journal.pcbi.1005944

92. Goel M, Sun H, Jiao W-B, Schneeberger K. SyRI: finding genomic rearrangements and local sequence differences from whole-genome assemblies. Genome Biol. 2019;20: 277. doi:10.1186/s13059-019-1911-0

93. Goel M, Schneeberger K. plotsr: visualizing structural similarities and rearrangements between multiple genomes. Bioinformatics. 2022;38: 2922–2926. doi:10.1093/bioinformatics/btac196

94. Darling ACE, Mau B, Blattner FR, Perna NT. Mauve: multiple alignment of conserved genomic sequence with rearrangements. Genome Res. 2004;14: 1394–1403. doi:10.1101/gr.2289704

95. Jiang T, Liu Y, Jiang Y, Li J, Gao Y, Cui Z, et al. Long-read-based human genomic structural variation detection with cuteSV. Genome Biol. 2020;21: 189. doi:10.1186/s13059-020-02107-y

